# Essential Gene Knockdowns Reveal Genetic Vulnerabilities and Antibiotic Sensitivities in *Acinetobacter baumannii*

**DOI:** 10.1101/2023.08.02.551708

**Authors:** Ryan D. Ward, Jennifer S. Tran, Amy B. Banta, Emily E. Bacon, Warren E. Rose, Jason M. Peters

## Abstract

The emergence of multidrug-resistant Gram-negative bacteria underscores the need to define genetic vulnerabilities that can be therapeutically exploited. The Gram-negative pathogen, *Acinetobacter baumannii*, is considered an urgent threat due to its propensity to evade antibiotic treatments. Essential cellular processes are the target of existing antibiotics and a likely source of new vulnerabilities. Although *A. baumannii* essential genes have been identified by transposon sequencing (Tn-seq), they have not been prioritized by sensitivity to knockdown or antibiotics. Here, we take a systems biology approach to comprehensively characterize *A. baumannii* essential genes using CRISPR interference (CRISPRi). We show that certain essential genes and pathways are acutely sensitive to knockdown, providing a set of vulnerable targets for future therapeutic investigation. Screening our CRISPRi library against last-resort antibiotics uncovered genes and pathways that modulate beta-lactam sensitivity, an unexpected link between NADH dehydrogenase activity and growth inhibition by polymyxins, and anticorrelated phenotypes that underpin synergy between polymyxins and rifamycins. Our study demonstrates the power of systematic genetic approaches to identify vulnerabilities in Gram-negative pathogens and uncovers antibiotic-essential gene interactions that better inform combination therapies.

**Importance:** *Acinetobacter baumannii* is a hospital-acquired pathogen that is resistant to many common antibiotic treatments. To combat resistant *A. baumannii* infections, we need to identify promising therapeutic targets and effective antibiotic combinations. In this study, we comprehensively characterize the genes and pathways that are critical for *A. baumannii* viability. We show that genes involved in aerobic metabolism are central to *A. baumannii* physiology and may represent appealing drug targets. We also find antibiotic-gene interactions that may impact the efficacy of carbapenems, rifamycins, and polymyxins, providing a new window into how these antibiotics function in mono- and combination therapies. Our studies offer a useful approach for characterizing interactions between drugs and essential genes in pathogens to inform future therapies.

## Introduction

The rise of antibiotic resistance in Gram-negative pathogens, including *Acinetobacter baumannii*, is a pressing healthcare concern, as many infections become untreatable amid a stalled pipeline for novel therapies (1). *A. baumannii* causes serious infections in hospitalized patients and is considered an urgent threat for its ability to evade killing by last-resort antibiotics (2). It has numerous defenses against antibiotics including a propensity to acquire resistance genes through horizontal transfer (3, 4), low membrane permeability coupled with robust efflux to prevent antibiotics from reaching their cytoplasmic targets (5), and rapid accumulation of resistance mutations (6). Although its unique strengths in resisting antibiotics are well documented, less is known about whether *A. baumannii* carries any unique vulnerabilities that could be therapeutically exploited.

The distinct physiology of *A. baumannii* sets it apart from well-studied, Gram-negative bacteria. Among the Gram-negative ESKAPE pathogens (*i.e.*, *Klebsiella*, *Acinetobacter*, *Pseudomonas*, and *Enterobacter*), *A. baumannii* is the only obligate aerobe, requiring oxidative phosphorylation to generate ATP (7). Further, the outer membrane of *A. baumannii* contains lipooligosaccharide (LOS) rather than lipopolysaccharide (LPS) found in most Gram-negative bacteria (8). LOS and LPS both contain a core lipid A moiety, but LOS lacks the repeating units of O-polysaccharide found in LPS (8). Although LPS is essential for viability in other Gram-negative ESKAPE pathogens, a recent study showed that LOS was dispensable in ∼60% of *A. baumannii* strains tested, including contemporary clinical isolates (9). LOS^-^ strains cannot be targeted by lipid A-binding antibiotics, such as polymyxins, increasing the antibiotic resistance threat posed by *A. baumannii* (10). Finally, *A. baumannii* has numerous genes of unknown function, including essential genes that are not present in model Gram-negatives or other ESKAPE pathogens (11). These distinctions underscore the importance of examining essential gene phenotypes and antibiotic interactions directly in *A. baumannii*.

Systematic genetic studies of *Acinetobacter* species have provided valuable physiological insights, although *A. baumannii* essential genes have not been comprehensively characterized. Tn-seq studies in *A. baumannii* identified putative essential genes (11, 12), defined phenotypes for previously uncharacterized genes (13), and uncovered the mechanism for strain-specific essentiality of LOS biosynthesis (9). An elegant Tn-seq study of non-pathogenic *Acinetobacter baylyi* monitored depletion of strains with disrupted essential genes following natural transformation (14), but it remains unclear whether those findings are directly applicable to *A. baumannii*.

CRISPR interference (CRISPRi) is the premier genetic tool to define essential gene function and antibiotic-gene interactions in bacteria. This gene knockdown technology uses a programmable, single guide RNA (sgRNA) to direct a catalytically inactive Cas effector protein (typically dCas9) to a target gene for silencing (15, 16). CRISPRi enables partial knockdown, and, thus, phenotyping of essential genes either by titrating the levels of CRISPRi components using inducible or weak promoters (15, 17, 18), or by modifying the sgRNA to weaken its interaction with target DNA (19–21) or dCas9 (22). Due to its portability, CRISPRi has proven valuable for phenotyping essential genes in diverse bacteria, including ESKAPE and other pathogens (11, 23, 24). Antibiotic-gene interaction screens using CRISPRi often recover the direct antibiotic target or related pathways among the largest outliers (17, 25). For instance, we previously identified the direct targets of two uncharacterized antibiotics using a *Bacillus subtilis* essential gene CRISPRi library, followed by genetic and biochemical validation of top hits (17). Although CRISPRi has been previously developed in *A. baumannii* by us and others (11, 23), only a handful of essential genes have been phenotyped to date.

To systematically probe for genetic vulnerabilities in *A. baumannii*, we generated and screened a pooled CRISPRi library targeting all putative essential genes [FIG 1A]. We identified essential genes and pathways that are most sensitive to knockdown, thereby prioritizing targets for future drug screens. We further used CRISPRi to define genetic interactions with last-resort antibiotics, finding antibiotic target pathways, obstacles to drug efficacy, and antibiotic-gene phenotypes that that may underlie synergistic drug combinations.

**Fig 1.**
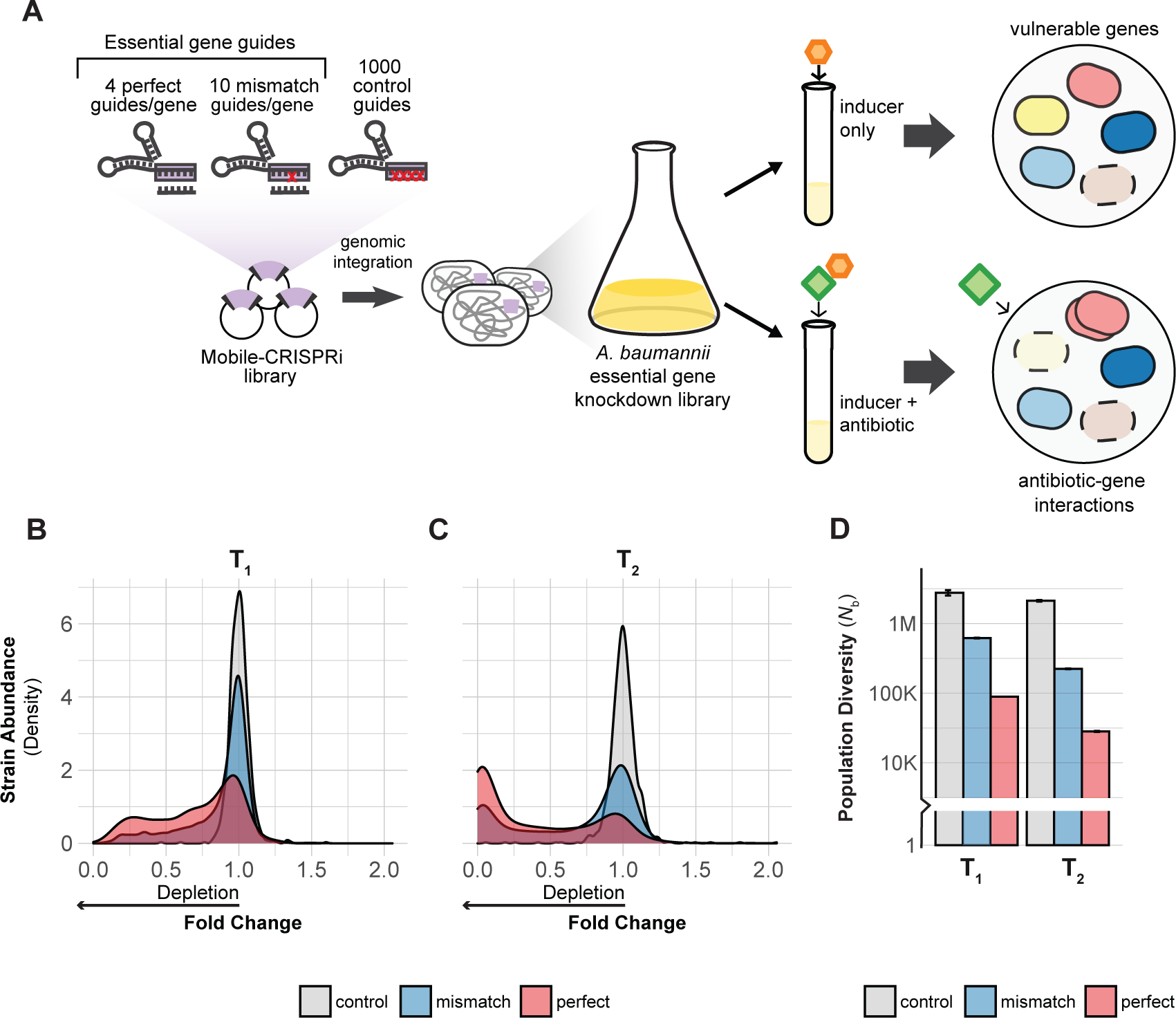
CRISPRi screening overview. **(A)** Design and construction of a Mobile-CRISPRi library targeting all putative essential genes in *A. baumannii* 19606. The library was screened with CRISPRi inducer (1 mM IPTG) to identify genes that are vulnerable to knockdown or with inducer and a sub-MIC concentration of antibiotic to identify antibiotic-gene interactions. **(B-C)** Density plot showing depletion of essential gene targeting sgRNA spacers (perfect match or mismatch) from the library but not depletion of non-targeting control sgRNAs during growth over two time points (T1 and T2) **(D)** The population diversity (*N_b_*) of essential gene targeting sgRNAs is reduced relative to controls, indicating that those sgRNAs are depleted during growth. The white horizontal line through the bars indicates a break in the data.

## Results

### Construction and validation of an *A. baumannii* essential gene CRISPRi library

We constructed a CRISPRi library targeting all putative essential genes in *A. baumannii* 19606, a strain extensively used to characterize the fundamental biology of *A. baumannii* that is also the type strain for antibiotic susceptibility testing (26). Notably, this strain is viable without LOS (9), allowing us to examine the phenotypic consequences of LOS loss. Developing our library in a susceptible strain also made it straightforward to use antibiotics as probes for gene function.

To systematically investigate essential genes in *A. baumannii,* we first optimized CRISPRi in *A. baumannii*, finding that reduced expression of *dcas9* lowered toxicity and still achieved ∼20-fold knockdown (Supplementary Methods, [FIG S1A-E, and Tables S1-4]). We next designed and constructed a CRISPRi library targeting all putative essential genes in *A. baumannii*. As the goal of our study was to characterize rather than define essential genes, we used existing Tn-seq data (12) to generate a list of CRISPRi targets we call the “*Ab* essentials” (406 genes total, [Table S5, S8]). We designed a computationally optimized CRISPRi library targeting the *Ab* essentials that consisted of three types of sgRNAs: 1) perfect match sgRNAs (15) to maximize knockdown (∼4/gene), 2) mismatch sgRNAs (19) to create a gradient of partial gene knockdowns (∼10/gene), and 3) control sgRNAs that are non-targeting (1000 total). This library was cloned and site-specifically integrated into the 19606 genome using Mobile-CRISPRi [FIG 1A] (23). Illumina sequencing of integrated sgRNA spacers confirmed that our CRISPRi library successfully targeted all the *Ab* essentials (median = 14 guides/gene; [FIG S2A]). Our approach, which includes using multiple sgRNAs per gene and robust statistics, mitigates potential issues with toxic or inactive guides.

To validate our *A. baumannii* CRISPRi library, we measured the depletion of essential-gene-targeting sgRNAs during pooled growth. We grew the library to exponential phase in rich medium (LB) without induction (T0), diluted back into fresh medium with saturating IPTG to induce CRISPRi and grew cells for ∼7 doublings (T1), then diluted back a second time in IPTG-containing medium and grew cells for an additional ∼7 doublings (T2). Quantifying strain depletion using log_2_ fold change (log_2_FC) and population diversity (*N*_b_; (27)) between T0, T1, and T2 ([FIG 1B-D, S2B], [Table S6]) revealed noticeable depletion of essential-gene-targeting sgRNAs by T1 and substantial depletion by T2, while control sgRNAs were unaffected. The lack of an induction effect on control strain abundance suggests that toxic guide RNAs such as “bad seeds” (28) are largely absent from our library. Taken together, our CRISPRi library effectively and comprehensively perturbs essential gene functions in *A. baumannii*.

### Identification of *A. baumannii* essential genes and pathways that are sensitive to knockdown

Essential genes with a strong, negative impact on fitness when knocked down, *i.e.*, “vulnerable” genes, are potential high-value targets for antibiotic development. CRISPRi enables the identification of vulnerable genes by controlling the duration and extent of knockdown (19, 20, 29). To define a set of vulnerable genes, we first quantified depletion of strains containing perfect match guides from the CRISPRi library during growth in rich medium (LB) [FIG S3A]. At T1, 88 genes showed significant depletion (log_2_FC<-1 and Stouffer’s p<0.05), and by T2 an additional 192 genes were depleted (280/406 total or 69%; Table S6). Screening our library in antibiotics at sub-MIC (minimal inhibitory concentration) levels recovered phenotypes for 74 for the 126 genes that were non-responsive in rich medium (see below), suggesting that these genes could be involved in antibiotic mode of action [FIG S3B]. The remaining 52 genes that were non-responsive in all our conditions may require >20-fold depletion (19), are false positives from the Tn-seq analysis used to define the *Ab* essentials (12), or are not essential in 19606. Overall, most *Ab* essentials (354/406 or 87%) showed significant phenotypes in our CRISPRi screens.

We sought to prioritize target genes and pathways by sensitivity to knockdown. Because CRISPRi knockdown affects transcription units (TUs) that can encode multiple gene products, we assigned essential genes to TUs and then organized the TUs into two groups: those containing only one essential gene and those containing multiple essential genes [Table S7]. We observed that most essential genes are in TUs containing either only one essential gene or multiple genes that participate in the same cellular process, limiting the phenotypic ambiguity of CRISPRi. Next, we ranked TU sensitivity to knockdown by the median log_2_FC of perfect match guides targeting essential genes present in the TU [Table S6]. Our measurements of log_2_FC are robust; however, we caution that small quantitative differences in gene/TU ranks may not always indicate meaningful variations in vulnerability.

Knockdowns of *murA*, *rpmB*, *aroC* and the poorly characterized gene, GO593_00515 were among the most depleted strains in our CRISPRi library [FIG 2A-B]. These genes represent established as well as underexplored therapeutic targets, and are in TUs containing only one essential gene, allowing straightforward interpretation of phenotypes. The *murA* gene, which encodes the target of fosfomycin (30), is vulnerable to knockdown despite fosfomycin’s inefficacy against *A. baumannii* due to efflux by the AbaF pump (31). L28, encoded by *rpmB*, is a bacterium-specific ribosomal protein that is required for assembly of the 70S ribosome in *Escherichia coli* (32, 33), but has no characterized inhibitors to our knowledge. Interestingly, *E. coli* cells with reduced L28 levels accumulate ribosome fragments that can be assembled into translation-competent ribosomes by expressing additional L28 (33), suggesting that L28 could play a role in regulation of ribosome assembly. The *aroC* gene encodes chorismate synthase, a metabolic enzyme genetically upstream of aromatic amino acid and folate biosynthesis. The abundance of aromatic amino acids in LB medium used in our screen suggests that the essential role of *aroC* is likely in folate biosynthesis. Chorismate synthase is essential in several bacterial species including Gram-positives, such as *B. subtilis* (34), and is vulnerable to knockdown in *Mycobacterium tuberculosis* (29), raising the possibility that *aroC* could be a general, high-value target.

**Fig 2.**
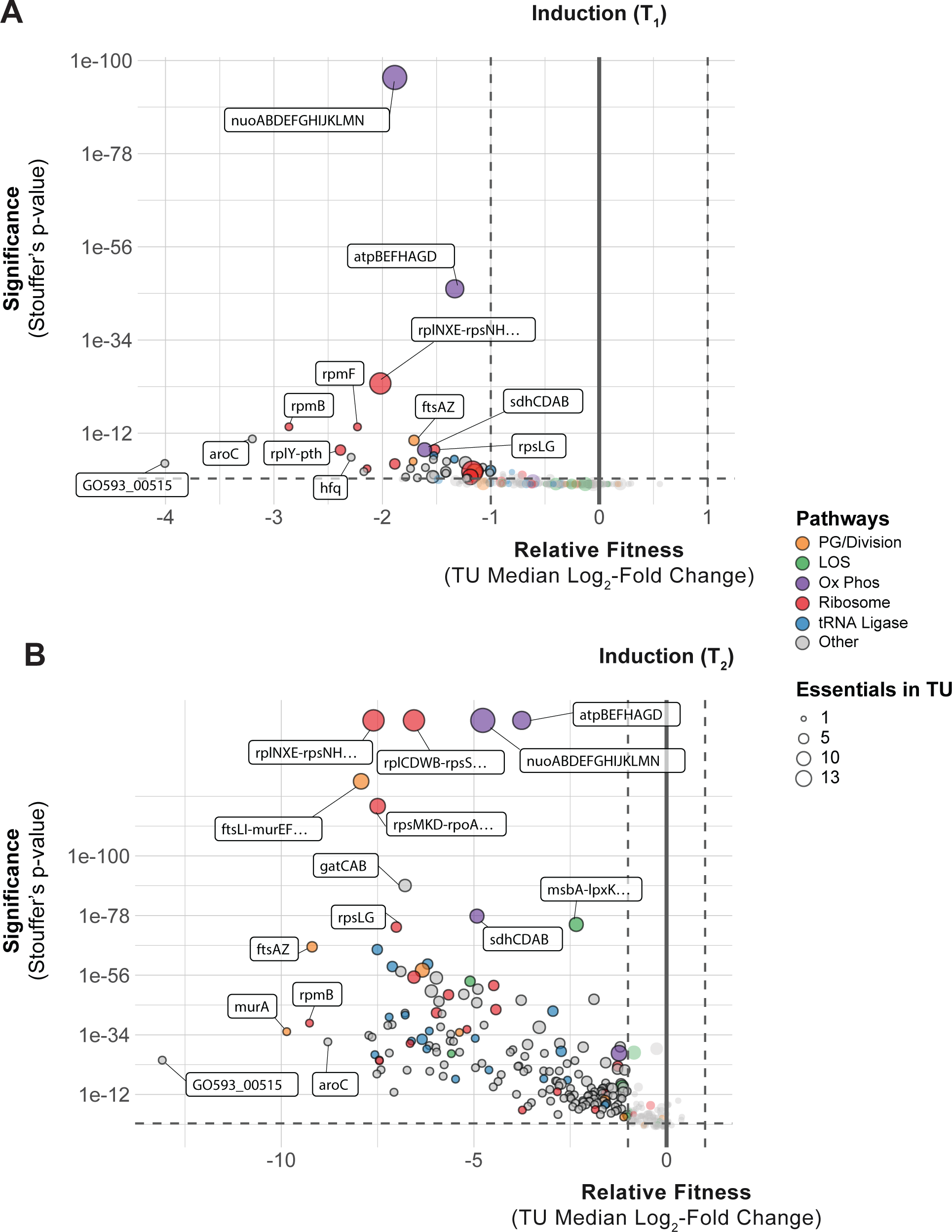
*A. baumannii* genes and pathways that are vulnerable to knockdown. **(A-B)** Depletion of sgRNAs targeting transcription units (TUs) from the CRISPRi library during growth in inducer over two time points (T_1_ and T2). Vertical dashed lines indicate a two-fold loss in fitness relative to non-targeting sgRNAs and horizontal dashed lines indicate a Stouffer’s *p* value of ≤ 0.05. Stouffer’s p values were calculated at the TU level by combining the false discovery rates (FDRs) of all individual sgRNAs targeting the TU. TUs related to pathways discussed in the text are colored according to the figure legend and the number of essential genes in a TU is indicated by point size.

Surprisingly, the most depleted knockdown strain in our library targeted an uncharacterized gene: GO593_00515 [FIG 2A-B]. GO593_00515 is predicted to encode an Arc family transcriptional repressor; Arc repressors have been extensively studied for their role in the Phage P22 life cycle (35). Accordingly, GO593_00515 is located within a predicted prophage in the 19606 genome; this locus is occupied by a similar but distinct prophage in the model resistant strain AB5075 [FIG S4A]. Synteny between the 19606 prophage and P22 suggested a role for GO593_00515 in lysogeny maintenance. Consistent with this hypothesis, we found that GO593_00515 knockdown cells showed little growth 10 hours after dilution into IPTG-containing medium [FIG S4C], and addition of IPTG to growing GO593_00515 knockdown cells caused complete lysis occurred within 7 hours [FIG S4B]. We reasoned that if the essential function of GO593_00515 is to repress expression of toxic prophage genes, we could suppress its essentiality by deleting the surrounding prophage genes entirely. Indeed, we recovered prophage deletion strains lacking GO593_00515 after inducing GO593_00515 in the presence of an integrated knockout plasmid [FIG S4C]. Thus, repression of toxic prophage genes is a critical but conditionally essential function in *A. baumannii*. Given the ubiquity of prophages harboring toxic lysis genes (36), we suggest that knockdown of phage repressors could aid in identifying proteins that are exceptional at lysing *A. baumannii*.

Sensitivity to knockdown among groups of genes with related functions provided further insight into *A. baumannii* vulnerabilities. Strong depletion of knockdowns targeting components of the ribosome, peptidoglycan (PG) synthesis, and cell division validated our CRISPRi screen by identifying pathways targeted by clinically relevant antibiotics [FIG 2A-B]. Genes encoding aminoacyl-tRNA synthetases (aaRSs) were functionally enriched among strains with reduced abundance at T2. Mupirocin, which targets IleRS, is the only inhibitor of a bacterial aaRS used clinically, although other aaRS inhibitors are used to treat infections caused by eukaryotic microbes [PMID: 33799176]. aaRSs are currently prioritized as targets for tuberculosis treatment as *M. tuberculosis* aaRS genes are vulnerable to knockdown (29) and a LeuRS/MetRS dual inhibitor is currently undergoing clinical trials (37). Our data demonstrate the vulnerability of aaRS genes in *A. baumannii* and suggest that aaRSs could serve as effective targets. Oxidative phosphorylation (oxphos) genes also stood out by degree of functional depletion in our library as early as T1 [FIG 2A]. Among the oxphos outliers, genes encoding the NADH dehydrogenase complex I (NDH-1; *nuo* genes) were particularly sensitive to knockdown. This finding highlights the distinct importance of aerobic metabolism in *A. baumannii* compared to other Gram-negative pathogens, such as *E. coli*, where NDH-1 is not essential for viability in aerobic conditions (38).

Ideal antibiotic targets have a tight relationship between target function and fitness such that small perturbations result in a substantial loss of viability. Recent work in model bacteria (19, 20) and *M. tuberculosis* (29) has found that the relationship between knockdown and fitness for essential genes is non-linear and varies by gene or pathway. To examine this phenomenon for *A. baumannii* vulnerable genes, we fit the relationship between gene knockdown (predicted by machine learning (19)) and fitness (log_2_FC of mismatch guides) to generate “knockdown-response” curves [FIG 3A-B]. We found that vulnerable genes and pathways were highly sensitive to even low levels of knockdown. Knockdown-response curves allowed us to determine the amount of knockdown required to elicit a half-maximal reduction in fitness (effective knockdown, or EK_50_) at the gene level. Vulnerable essential genes, such as *murA*, showed a substantial fitness defect at less than half of the maximal knockdown, whereas non-essential genes, such as *lpxA*, showed little fitness defects even at higher levels of knockdown. Other vulnerable genes (e.g., *rpmB*, *aroC*, and GO593_00515) also showed heightened sensitivity to knockdown [Fig. FIG S5]. We extended our knockdown-response analysis to the pathway level, finding that pathways with many vulnerable genes (PG/division) required less knockdown, on average, than pathways with few vulnerable genes (LOS) [FIG 3C-D]. Interestingly, although fitness at T2 was generally lower than T1 for vulnerable genes, EK_50_values at both time points were statistically indistinguishable. This demonstrates that even guides with weak knockdown contribute to vulnerability phenotypes and suggests that gene phenotypes require a certain threshold of knockdown that is pathway dependent in *A. baumannii*.

**Fig 3.**
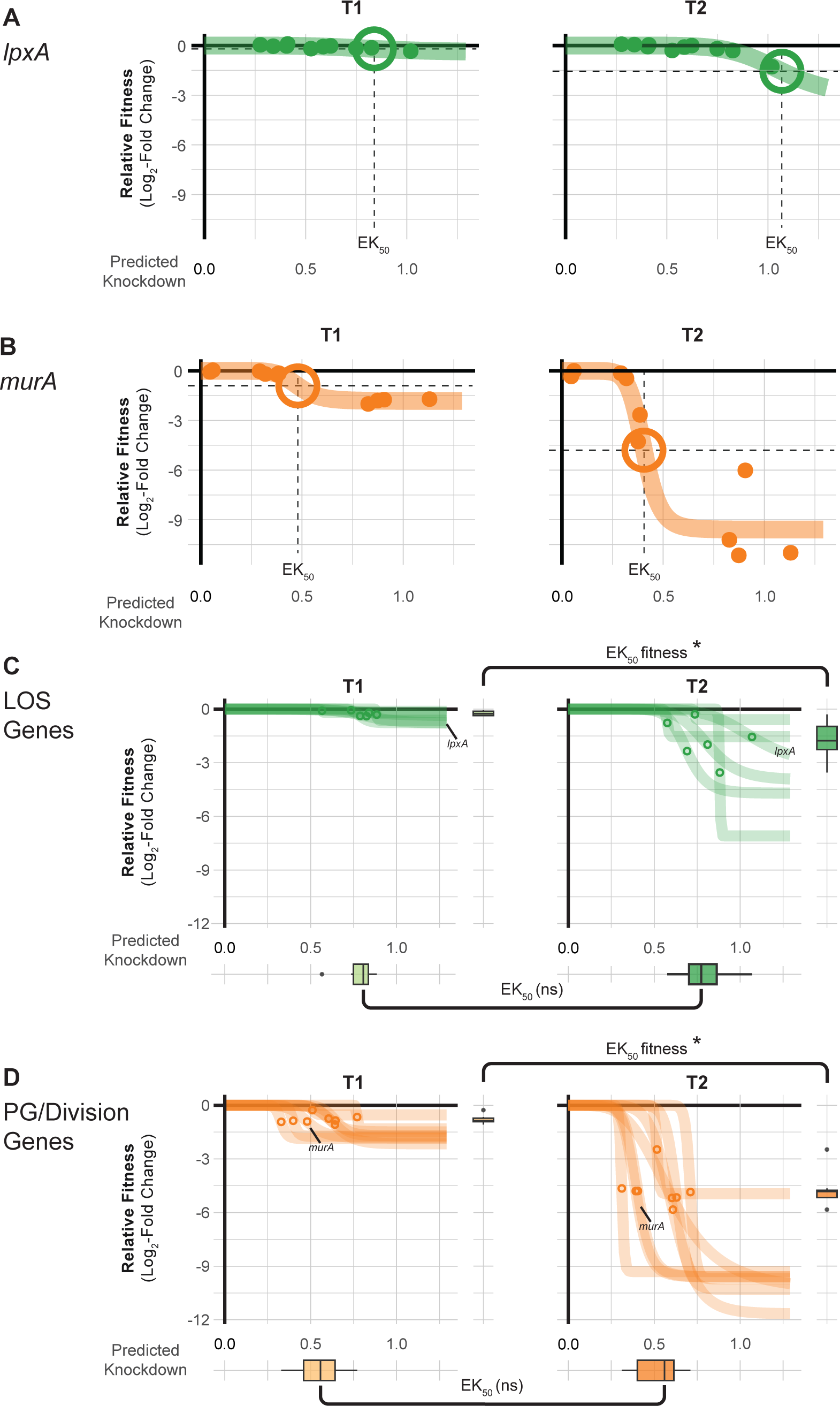
Knockdown-response curves describe gene and pathway vulnerability. **(A-B)** Knockdown-response curves of the LOS gene *lpxA* and the PG/division gene *murA*. Points are individual mismatch sgRNAs; mismatch sgRNA knockdown was predicted as previously described (19). Colored lines are a 4-parameter logistic fit describing the relationship between relative fitness and knockdown. The effective knockdown 50 (EK_50_) is the amount of predicted knockdown needed to achieve a half-maximal effect on relative fitness. EK_50_s are depicted as crosshairs. **(C-D)** Knockdown response curves for genes in LOS synthesis or PG/division pathways. Points indicate the EK_50_ for individual pathway genes. Boxplots on the y-axis show the distribution of relative fitness at EK_50_ for genes in the pathway and boxplots on the x-axis show the distribution of EK_50_ values for genes in the pathway. Statistical significance was assessed using Wilcoxon Rank Sum Test; asterisks indicate *p* ≤ 0.05 and ns for not significant.

### Essential gene knockdowns that potentiate or mitigate carbapenem sensitivity in *A. baumannii*

Antibiotic-gene interaction screens have the potential to identify targets that synergize with or antagonize existing therapies. Carbapenems, a class of beta-lactam antibiotics, are first-line treatments for *A. baumannii* that block PG synthesis by inhibiting penicillin binding proteins (PBPs) (39). To uncover antibiotic-essential gene interactions that impact sensitivity to carbapenems, we screened our CRISPRi library against sub-MIC concentrations of imipenem (IMI) and meropenem (MER) [FIG 4A, S6A-B]. We found that knockdown of genes involved in cell wall synthesis, including the direct target (*ftsI*, TU: *ftsLI-murEF-mraY*), increased carbapenem sensitivity. Knockdowns of genes required for PG precursor synthesis (*murA*, *dapA*) and translocation (*murJ*) were strongly depleted in both IMI and MER. MurA catalyzes the first committed step of PG synthesis, DapA is part of a pathway that converts L-aspartate to meso-diaminopimelate which is incorporated into PG precursors by MurE, and MurJ, the lipid II flippase, translocates PG precursors from the inside to the outside of the cytoplasmic membrane (40). To validate screen hits, we developed a high-sensitivity assay that uses Next Generation Sequencing to measure competitive fitness between a non-targeting and CRISPRi knockdown strain we call “CoMBaT-seq” (<u>Co</u>mpetition of <u>M</u>ultiplexed <u>Ba</u>rcodes over <u>T</u>ime).

**Fig 4.**
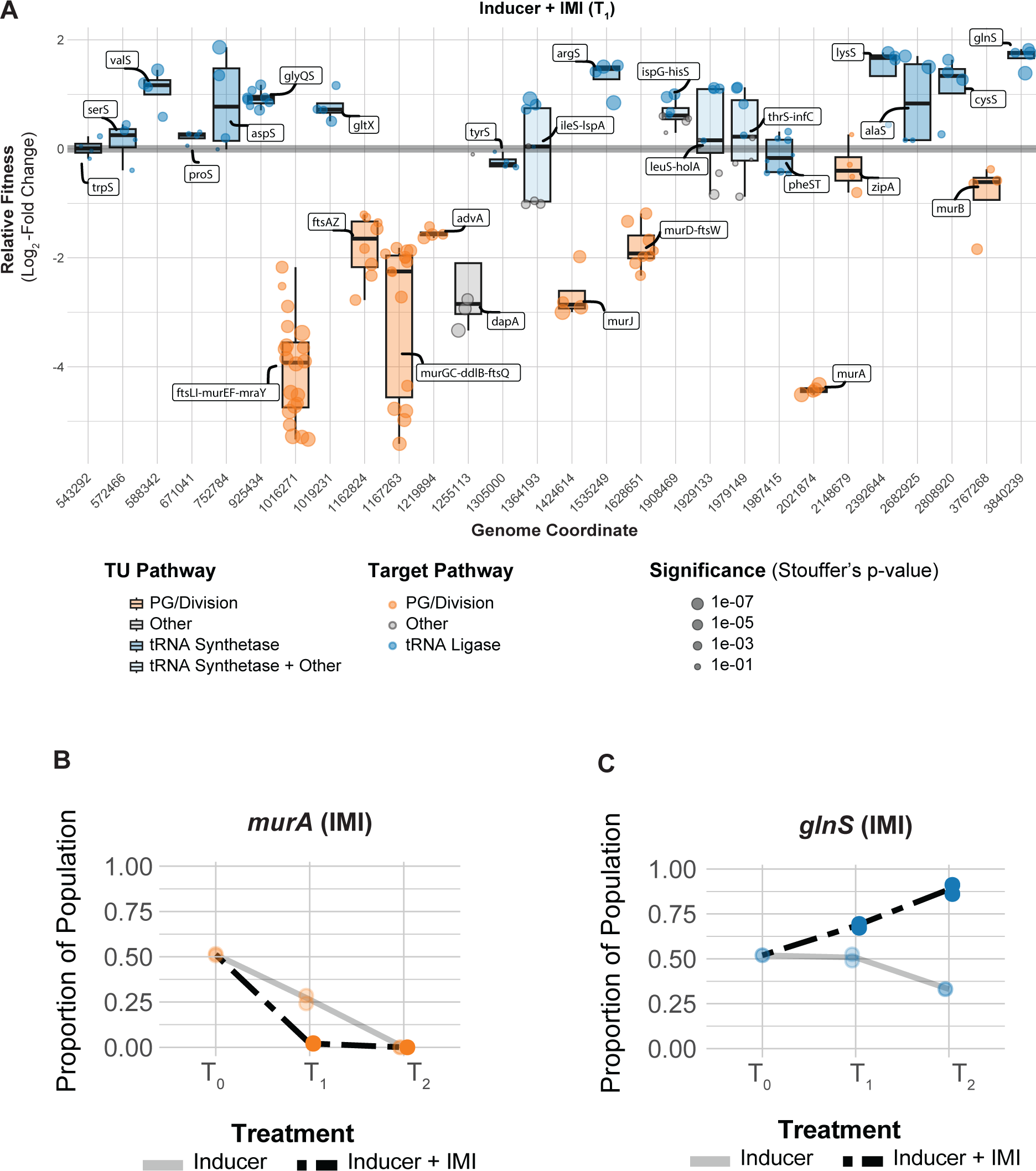
Essential gene interactions with carbapenem antibiotics in *A. baumannii*. **(A)** Boxplots showing the relative fitness of selected TUs that interact with imipenem (IMI) across the genome at T1. Points are individual genes in the TU. Boxplots are colored by relevant pathways; light-blue boxplots indicate TUs where tRNA synthetase genes are present with genes in other pathways. **(B-C)** CoMBaT-seq data from a growth competition between either a *murA* or *glnS* knockdown strain and a non-targeting control strain in the presence or absence of IMI. Only data from the gene targeting strain is depicted as the non-targeting control is the remaining proportion of the population. Points are data from individual experiments (N = 2).

CoMBaT-seq recapitulated *murA* vulnerability to knockdown and further sensitivity to IMI [FIG 4B]. Consistent with our *murA*-IMI interaction, we found that fosfomycin and IMI synergize in *A. baumannii* [FIG S7A], as is the case in other Gram-negative pathogens (e.g., *Pseudomonas aeruginosa* (41)). Although no clinically relevant inhibitors of DapA and MurJ exist, to our knowledge, we speculate that such inhibitors would have the potential to synergize with carbapenems. Intriguingly, knockdowns of *advA*—an Acinetobacter-specific division gene (13)— were also sensitized to carbapenems, raising the possibility of *A. baumannii* targeting combination therapies should inhibitors of AdvA be identified.

Gene knockdowns that mitigate antibiotic function can reveal routes to resistance or target combinations that result in antagonism and should be avoided therapeutically. Given that increasing carbapenem resistance is an urgent clinical concern for *A. baumannii*, we sought to identify genes and pathways that mitigate the efficacy of IMI and MER. Although previous work suggested that growth rate and beta-lactam resistance are linearly related (42, 43), we found only a modest linear relationship growth and IMI/MER resistance across knockdown strains in our library (R2 = 0.005, and 0.007, respectively [FIG S8A-B]). This indicates that slow growing strains of *A. baumannii* are not necessarily more resistant to beta-lactam treatment. Instead, we found that specific genetic pathways govern carbapenem resistance. Using gene set enrichment analysis, we identified ribosomal protein genes as a pathway that increases resistance to IMI/MER when perturbed [**IMI**: enrichment score = 4.65, FDR (afc) = 2.12e-06; **MER**: enrichment score = 2.43, FDR (afc) = 0.002], consistent with antagonism between beta-lactams and ribosome inhibitors described for other bacteria (44).

aaRS genes also emerged from our enrichment analysis [**IMI**: enrichment score = 4.93, FDR (afc) = 1.04e-06; **MER**: enrichment score = 5.04, FDR (afc) = 1.16e-06], uncovering a connection between tRNA charging and carbapenem resistance, as well as a surprising relationship between knockdown and fitness unique to antagonistic interactions. In particular, a subset of aaRS gene knockdowns including *argS*, *lysS*, *valS*, *cysS* and glnS showed increased relative fitness our IMI pooled screen [FIG 4A, S8C]. Although *glnS* resistance to IMI in MIC test strip and growth curve assays was modest [FIG S9], our more sensitive CoMBaT-seq assay showed a clear growth advantage for the *glnS* knockdown when competed against a non-targeting control in contrast to sensitive knockdowns such as *murA* [FIG 4B-C]. Our observations that the *glnS* knockdown is depleted during growth in rich medium and enriched during growth in IMI implied that the relationship between knockdown and fitness changed across the two conditions. Indeed, a 4-parameter knockdown-response curve fit well to mismatch guides targeting *glnS* without treatment, but poorly to the same guides in IMI treatment [FIG 5A-D, S10A]. Remarkably, IMI treated *glnS* knockdown strains showed increased relative fitness as knockdown increased up until a point at the strains lost viability, presumably due to a lack of glutamine tRNA charging. This pattern is reminiscent of a hormetic response in dose-response curves (45) where a low amount of drug produces a positive response that eventually becomes negative at higher doses [FIG 5A-B]. Accordingly, a 5-parameter logistic curve typically used in the context of hormetic responses improved the fit to IMI treated *glnS* mismatch strains but did not improve the fit of untreated strains [FIG 5C-D, S10B]. To test if the hormetic effect we observed between IMI and *glnS* in an antibiotic-gene interaction was relevant to antibiotic-antibiotic interactions, we measured the growth of wild-type *A. baumannii* treated with IMI and the aaRS inhibitor, mupirocin. Consistent with hormesis, IMI antagonized the effect of mupirocin at low concentrations, but had no positive impact on growth at higher concentrations [FIG 5E]. Although mupirocin treatment is not clinically relevant for *A. baumannii* due to high-level resistance, our work provides a proof of principle that hormetic effects can be predicted by genetic approaches and influence antibiotic susceptibility.

**Fig 5.**
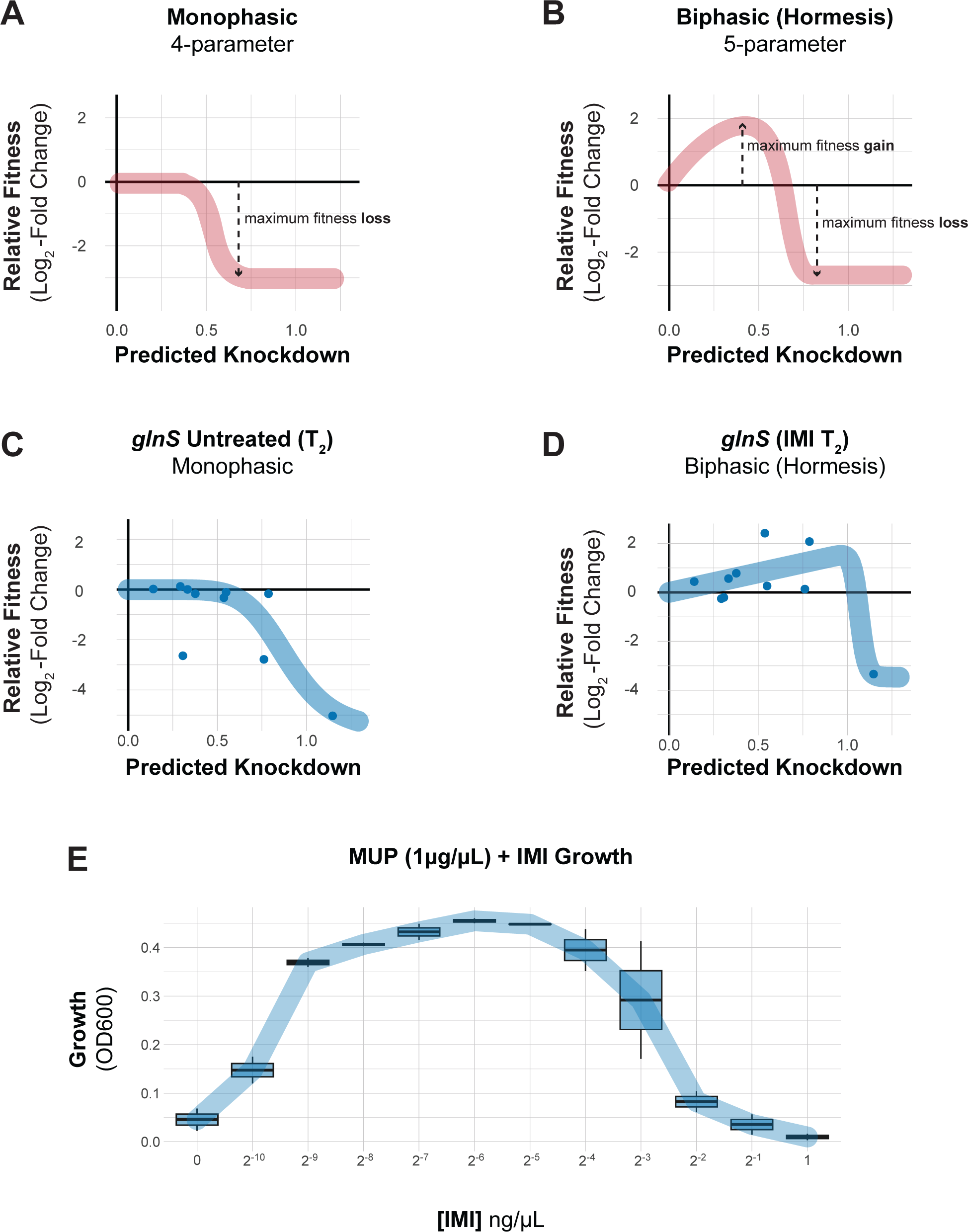
Knockdown extent affects the sign of antibiotic-gene interactions. **(A-B)** Schematics of idealized dose-response curves showing monotonic or hormetic relationships between dose and response; hormetic responses change the sign of the response depending on dose. **(C-D)** Knock-down-response curves of *glnS* show a nearly monotonic response in the absence of IMI, but a hormetic response in the presence of IMI. **(E)** The interaction between the IleRS tRNA synthetase inhibitor mupirocin (MUP) and IMI shows a hormetic response at intermediate concentrations of IMI.

### Anticorrelated phenotypes underlie synergy between colistin and rifampicin

Antibiotic-gene interaction screens can identify genes and pathways that underpin synergistic drug combinations, clarifying the genetic basis for synergy. Colistin (COL) and rifampicin (RIF) synergistically inhibit *A. baumannii* growth by an unknown mechanism [FIG S11](46). To define antibiotic-gene interactions that may inform COL-RIF synergy, we screened our CRISPRi library against COL and RIF individually. We found strong, opposing phenotypes in COL and RIF for genes encoding NDH-1 and LOS biosynthesis genes. COL, a polymyxin class antibiotic, is a last-resort treatment for carbapenem-resistant *A. baumannii* (47). COL binds to the lipid A moiety of LOS and is thought to kill cells by membrane disruption (48); complete loss of LOS results in a >500-fold increase in COL resistance (9). As expected, screening our library against a sub-MIC dose of COL identified LOS synthesis genes as resistant outliers [FIG 6, S12]. Among the most resistant outliers were *lpxC* (TU: *lpxC*) and *lpxA* (TU: *lpxD*-*fabZ*-*lpxA*), which encode enzymes that catalyze the first two committed steps in LOS synthesis and are commonly found in selections for COL resistant mutants (9). Genes involved in fatty acid biosynthesis biosynthesis (TU: *fabDG*, TU: *aroQ*-*accBC*) also showed increased resistance to COL, possibly by limiting the pool of fatty acids available for LOS synthesis [FIG S12]. Surprisingly, knockdown of genes encoding NDH-1 (TU: *nuoABDEFGHIJKLMN*) caused heightened sensitivity to COL in the context of our pooled screen [FIG 6, S9]. We robustly confirmed the COL sensitivity of a *nuoB* knockdown using our CoMBaT-seq assay [FIG 7A], although MIC test strips showed a more muted effect [FIG S13]. NDH-1 couples conversion of NADH to NAD+ to proton translocation across the inner membrane, but whether the key role for NDH-1 in *A. baumannii* physiology is NAD+ recycling or contributing to membrane potential (Δψ) is unknown. To address this issue, we measured the NAD+/NADH ratio and Δψ using an enzyme-coupled luminescence assay (NAD/NADH-Glo) and the membrane potential-sensitive dye Thioflavin T (ThT), respectively [FIG 7B, S14A]. Knockdown of *nuoB* lowered the NAD+/NADH ratio, consistent with reduced conversion of NADH to NAD+ by NDH-1 [FIG 7B].

**Fig 6.**
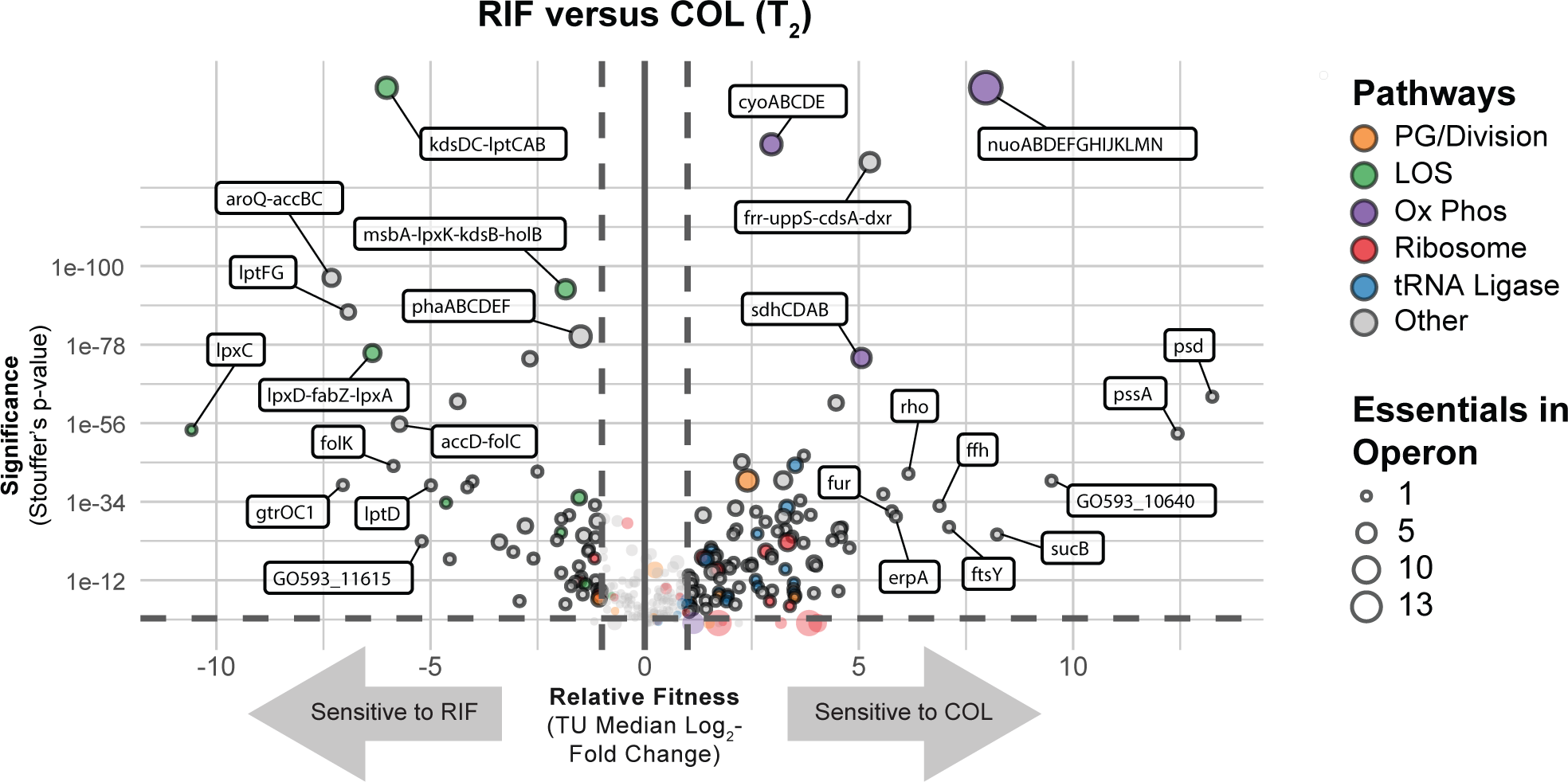
Essential gene knockdown phenotypes in rifampicin (RIF) versus colistin (COL). Depletion of sgRNAs targeting transcription units (TUs) from the CRISPRi library during growth in inducer and RIF or COL at T2. Vertical dashed lines indicate a two-fold loss in fitness relative to non-targeting sgRNAs and horizontal dashed lines indicate a Stouffer’s *p* value of ≤ 0.05. Stouffer’s p values were calculated at the TU level by combining the false discovery rates (FDRs) of all individual sgRNAs targeting the TU. TUs related to pathways discussed in the text are colored according to the figure legend and the number of essential genes in a TU is indicated by point size.

**Fig 7.**
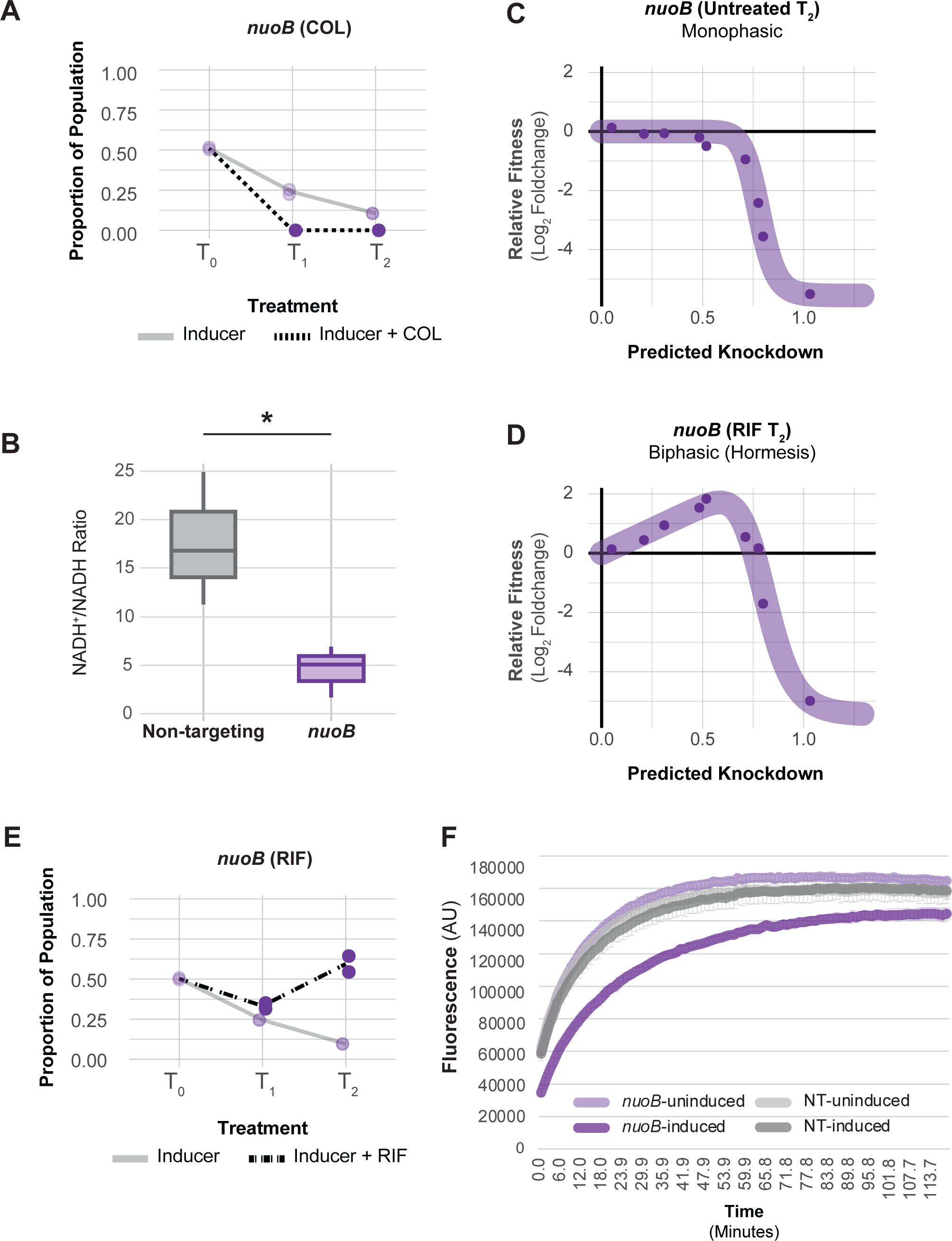
COL/RIF interaction with and physiological characterization of NDH-1 knockdown. **(A)** CoMBaT-seq data from a growth competition between a *nuoB* knockdown strain and a non-targeting control strain in the presence or absence of COL. Only data from the gene targeting strain is depicted as the non-targeting control is the remaining proportion of the population. Points are data from individual experiments (N = 2). **(B)** Measurement of the NAD+/NADH ratio in *nuoB* knock-down and non-targeting cells using the NAD/NADH-Glo assay. An unequal variance t-test was performed and the asterisk indicates that the p value ≤ 0.05. **(C-D)** Knockdown-response curves of *nuoB* show a nearly monotonic response in the absence of RIF, but a hormetic response in the presence of RIF. **(E)** CoMBaT-seq data from a growth competition between a *nuoB* knockdown strain and a non-targeting control strain in the presence or absence of RIF. **(F)** EtBr permeability assay of non-targeting and *nuoB* knockdown strains; *nuoB* knockdowns show decreased access of EtBr to DNA in the cytoplasm.

Unexpectedly, *nuoB* knockdown did not impact Δψ, although reduced Δψ in cells treated with the ionophore CCCP was readily apparent in our ThT assay [FIG S14B]. Thus, recycling of NADH to NAD+ for use in the TCA cycle, rather than maintenance of membrane potential, may be the critical cellular role of NDH-1. *A. baumannii* also encodes a non-essential, non-proton pumping NDH-2 enzyme that can be inhibited by COL *in vitro*. We speculate that NDH-2 inhibition by COL combined with knockdown of NDH-1 critically reduces cellular NAD+ levels, leading to enhanced sensitivity.

Rifampicin is a relatively large antibiotic (822.9 Da) that targets RNA polymerase (RNAP) in the cytoplasm but is typically avoided for treating Gram-negative infections due to low permeability (49). Consistent with a permeability barrier to rifampicin function (50), we found that knockdown of LOS synthesis and transport genes strongly sensitized cells to rifampicin [enrichment score = 8.26, FDR (afc) = 3.97e-05]. Again, knockdown of genes encoding NDH-1 produced an unexpected phenotype, this time increasing RIF resistance by an unknown mechanism [FIG 6, S15]. To further characterize the NDH-1 RIF resistance phenotype, we examined the knockdown-response curve of *nuoB* with and without RIF treatment. As seen previously with *glnS*, *nuoB* knockdown showed a hormetic response: increasing knockdown of *nuoB* increased relative fitness in RIF until the highest levels of *nuoB* knockdown where growth decreased [FIG 7C-D, S10C-D]. Although MIC changes were modest [FIG S16], our CoMBaT-seq assay showed a clear fitness benefit for *nuoB* knockdown in RIF relative to a non-targeting control [FIG 7E]. We considered that NDH-1 knockdown cells may have reduced permeability, limiting RIF entry into the cytoplasm. To test permeability, we measured uptake of ethidium bromide (EtBr) which fluoresces when bound to DNA in the cytoplasm [FIG 7F]. We found that *nuoB* knockdown cells had a reduced rate of EtBr uptake, demonstrating that cells with reduced NDH-1 activity are less permeable and suggesting a possible mechanism for increased RIF resistance.

COL and RIF showed strong, anticorrelated phenotypes in our CRISPRi screen [FIG 8], with LOS related knockdowns causing resistance to COL and sensitivity to RIF and NDH-1 knockdowns resulting in sensitivity to COL and resistance to RIF. These results support a collateral sensitivity model for COL-RIF synergy (51) (see Discussion). Taken together, COL and RIF synergistically kill *A. baumannii* by exerting opposite effects on LOS and NDH-1 that cannot be compensated for by reducing the function of either pathway.

**Fig 8.**
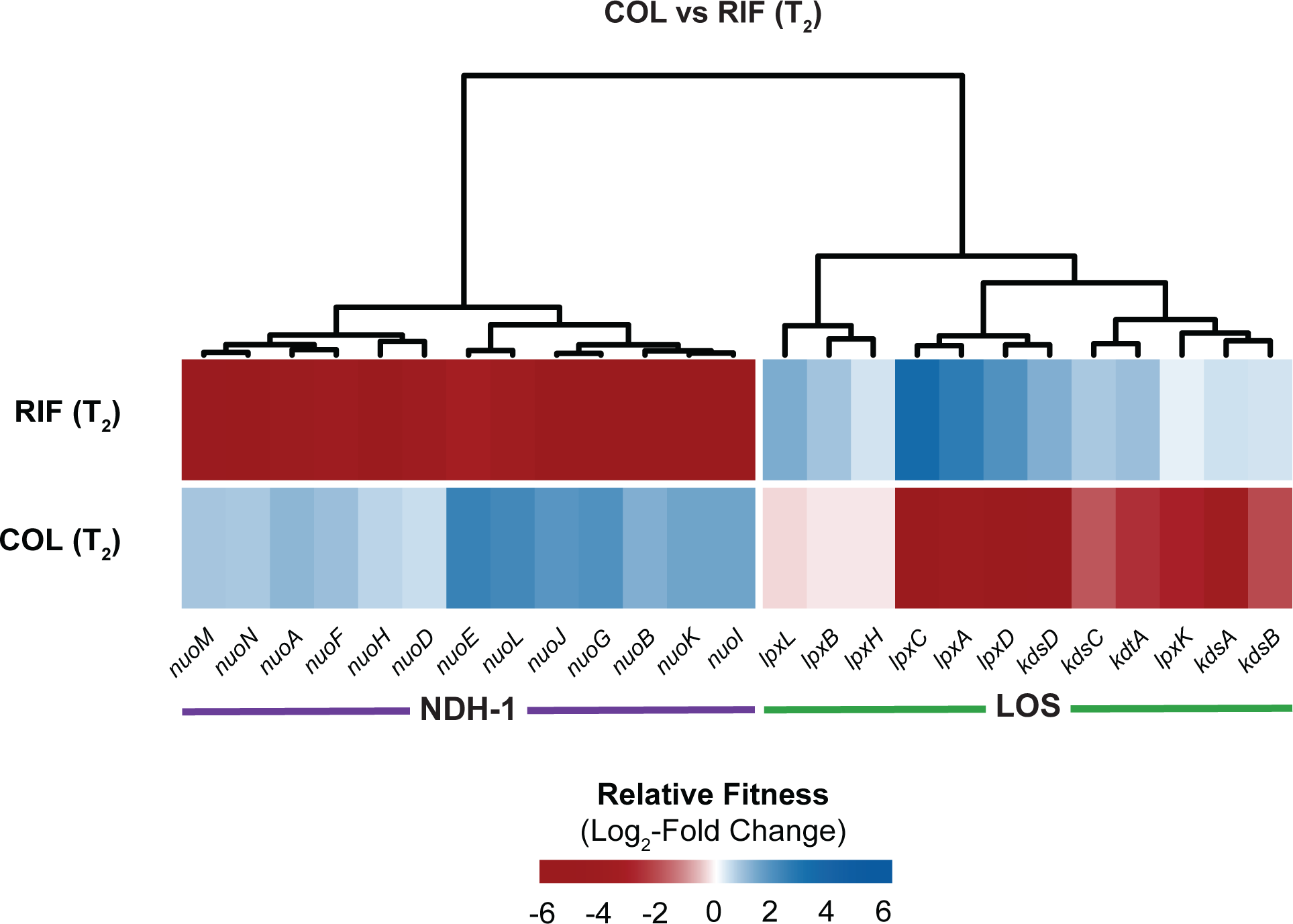
Anticorrelated gene-antibiotic interactions for colistin (COL) and rifampicin (RIF). Relative fitness changes for genes encoding NDH-1 or involved in LOS biosynthesis in COL or RIF treated conditions relative to untreated.

## Discussion

Bacterial susceptibility to antibiotics is underpinned by species- and condition-specific gene essentiality. The recent lack of innovative treatments for *A. baumannii* and other Gram-negative pathogens can be attributed to our limited knowledge of genetic weaknesses in these bacteria. This work advances our understanding of genetic vulnerabilities in *A. baumannii* by systematically perturbing and phenotyping essential genes. Using CRISPRi to knock down essential gene products, we identified genes that are sensitive to knockdown as well as genes that potentiate or mitigate antibiotic action. Together, these studies define potential targets for antibiotic discovery and provide a genetic approach for understanding synergistic therapies that is broadly applicable.

Our study of essential gene knockdown phenotypes in *A. baumannii* points to both unique and shared genetic vulnerabilities with other bacterial species. Our finding that *A. baumannii* is highly sensitive to depletion of genes encoding NDH-1 highlights a unique weakness in pathogens that are obligate aerobes and a possible therapeutic target. Among the Gram-negative ESKAPE pathogens, only *A. baumannii* is known to require *nuo* genes for aerobic growth in rich medium (52). Recent work from Manoil and colleagues in the non-pathogenic model strain, *Acinetobacter baylyi*, found that genes involved in oxidative phosphorylation were among the first to be depleted from a pool of transposon mutants (14); combining these observations with our CRISPRi results suggests that oxygen-dependent energy production is a physiological linchpin across the *Acinetobacter* genus. Our finding that *A. baumannii* genes involved in PG synthesis and translation are vulnerable to depletion underscores the conserved importance of these pathways across bacterial species (19, 29) and their foundational role as antibiotic targets.

Our finding that knockdown gradients of essential genes treated with antibiotics can mimic hormetic effects seen in dose-response curves (45) has implications for modeling conditional phenotypes of essential genes and dosing of combination therapies. For most essential genes, complete loss of gene function results in lethality under the majority of conditions. However, our mismatch guide strategy allowed us to examine intermediate levels of essential gene function that may be analogous to partial loss of function alleles found in resistant clinical isolates or adaptive evolution experiments. Partial loss of function mutants can exhibit striking differences in phenotype over a narrow range of function, as we observed with *glnS* and *nuoB* resistance during IMI and RIF treatment, respectively. These hormetic resistance phenotypes fit poorly to established 4-parameter logistic models, emphasizing the importance of considering alternative model parameters and comprehensive statistical approaches when quantifying intricate biological processes. Given our limited set of screening conditions, it is currently unclear how widespread the phenomenon of hormesis is for antibiotic-gene interactions, although we note that clear instances of hormesis were rare in our data. Hormesis in antibiotic interactions may have clinical relevance as well, as doses of combination therapies falling within the concentration window of a hormetic/antagonistic response would be ineffective. Our ability to predict antagonism between an aaRS inhibitor and carbapenems based on genetic data suggests that screening for antibiotic-gene interactions will have as much value in avoiding antagonisms as it does in identifying potential synergies.

Our data show an unexpected link between NADH dehydrogenase activity and growth inhibition by COL. NDH-1 knockdown strains were highly sensitized to COL in competitive growth assays, but the precise mechanism behind this sensitivity is unclear. Based on our measurements, NDH-1 knockdown primarily affects the ratio of NADH to NAD+ in cells, rather than membrane potential. COL inhibits conversion of NADH to NAD+ by the type II NADH dehydrogenase (NDH-2) in a purified system (53), although at much higher concentrations than used in our experiments. We speculate that the sensitivity of NDH-1 knockdowns to COL is due insufficient recycling of NAD+, which would be expected to reduce flux through the TCA cycle. In this scenario, CRISPRi knockdown reduces NDH-1 activity while COL inhibits NDH-2 activity, resulting in further skewing of the NADH/NAD+ ratio toward NADH and away from NAD+. Flux through the TCA cycle would be expected to decrease as multiple steps in the cycle require available NAD+ (54, 55). In general, identifying targets that potentiate COL activity may be clinically relevant in the context of combination therapy because toxicity is a major dose-limiting concern of polymyxin antibiotics (56). Employing effective combination treatments using colistin concentrations below toxicity thresholds would greatly improve its clinical utility and safety against *A. baumannii*. Our CRISPRi approach could inform not only combinations with polymyxins, but also other antibiotics which have dose-limiting toxicity concerns that prevent more widespread use.

COL and RIF have been shown to synergistically kill *A. baumannii* and other Gram-negatives (56) by an unknown mechanism. We suggest that the synergy can be explained by a collateral sensitivity mechanism (51), in which genetic perturbations that promote COL resistance increase sensitivity to RIF and *vice versa*. Treatment with COL selects for mutations in LOS biosynthesis genes (10), while the loss of LOS promotes permeability to RIF (and other antibiotics (56)). Accordingly, the presence of RIF has been shown to reduce recover of inactivated *lpx* genes in selections for COL resistance (57). Mutations in *nuo* genes are commonly obtained in screens for tobramycin resistance in *P. aeruginosa* (58, 59), supporting a model in which reduced NDH-1 function decreases permeability of the inner membrane to antibiotics. Consistent with this model, we found that EtBr fluorescence, which is often used as a proxy for measuring permeability of small molecules, was decreased in NDH-1 knockdown strains. Collateral sensitivity comes into play when mutations that reduce NDH-1 activity to block rifampicin entry increase sensitivity to colistin which further reduces NDH-1 function. This mechanism of synergy can impact other Gram-negative ESKAPE pathogens but is particularly relevant in *A. baumannii* because LOS is not essential and NDH-1 is uniquely required for viability. In the case of colistin and rifampicin, collateral sensitivity manifests as anticorrelated phenotypes in chemical genomics data. We speculate that anticorrelated phenotypic signatures are predictive of antibiotic synergy, particularly in the context of bacteria with low permeability such as *A. baumannii* and *P. aeruginosa*. Interrogating a larger chemical genomics dataset with a greater diversity of antibiotics for these organisms will shed light on general rules for antibiotic-gene interactions and their implications for discovering synergy.

## Materials and Methods

### Strains and growth conditions

Strains are listed in Table S1. Details of strain growth conditions are described in the Supporting Information.

### General molecular biology techniques and plasmid construction

Plasmids and construction details are listed in Table S2. Oligonucleotides are listed in Table S3. Details of molecular biology techniques are described in the Supporting Information.

### *A. baumannii* Mobile-CRISPRi system construction

An *A. baumannii* strain with the Mobile-CRISPRi (MCi) system from pJMP1183 (23) inserted into the *att*_Tn*7*_ site (Fig. S1A), which constitutively expresses mRFP and has an mRFP-targeting sgRNA, has a growth defect when induced with 1mM IPTG (Fig. S1B; “parent”). Strains with suppressors of the growth defect that still maintained a functional CRISPRi system were identified by plating on LB supplemented with 1mM IPTG and selecting white colonies (red colonies would indicate a no longer functional MCi system; Fig. S1B and S1C). gDNA was extracted and mutations in the dCas9 promoter were identified by Sanger sequencing (Fig. S1D). The Mobile-CRISPRi plasmid pJMP2748 is a variant of pJMP2754 (Addgene 160666) with the sgRNA promoter derived from pJMP2367 (Addgene 160076) and the dCas9 promoter region amplified from the *A. baumannii* suppressor strain gDNA with oJMP635 and oJMP636. Plasmid pJMP2776, which was used to construct the *A. baumannii* essential gene library and individual sgRNA constructs, was created by removal of the GFP expression cassette from pJMP2748 by digestion with PmeI and re-ligation. This system shows ∼20-fold knockdown when targeting the *GFP* gene (Fig. S1E). Plasmids will be submitted to Addgene.

### *A. baumannii* Mobile-CRISPRi individual gene and gene library construction

sgRNAs were designed to knockdown essential genes in *A. baumannii* 19606 using a custom python script and Genbank accession #s CP046654.1 and CP046655.1 as detailed in reference (60). Mismatch guides were designed and predicted knockdown was assigned as previously described (19). sgRNA-encoding sequences were cloned between the BsaI sites of Mobile-CRISPRi (MCi) plasmid pJMP2776. Methodology for cloning individual guides was described previously in detail (60). Briefly, two 24-nucleotide (nt) oligonucleotides encoding an sgRNA were designed to overlap such that when annealed, their ends would be complementary to the BsaI-cut ends on the vector.

The pooled essential gene CRISPRi library was constructed by amplification of sgRNA-encoding spacer sequences from a pooled oligonucleotide library followed by ligation into the BsaI-digested MCi plasmid. Specifically, a pool of sgRNA-encoding inserts was generated by PCR amplification with primers oJMP697 and oJMP698 from a 78-nt custom oligonucleotide library (2020-OL-J, Agilent) with the following conditions per 500 µl reaction: 100 µl Q5 buffer, 15 µl GC enhancer, 10 µl 10mM each dNTPs, 25 µl each 10 µM primers oJMP897 and oJMP898, 10 µl 10 nM oligonucleotide library, 5 µl Q5 DNA polymerase, and 310 µl H2O with the following thermocycling parameters: 98°C, 30s; 15 cycles of: 98°C, 15s; 56°C, 15s; 72°C, 15s; 72°C, 10 min; 10°C, hold. Spin-purified PCR products were digested with BsaI-HF-v2 (R3733; NEB) and the size and integrity of full length and digested PCR products were confirmed on a 4% agarose e-gel (Thermo). The BsaI-digested PCR product (without further purification) was ligated into a BsaI-digested MCi plasmid as detailed in (60). The ligation was purified by spot dialysis on a nitrocellulose filter (Millipore VSWP02500) against 0.1 mM Tris, pH 8 buffer for 20 min prior to transformation by electroporation into *E. coli* strain BW25141 (sJMP3053). Cells were plated at a density of ∼50,000 cells/plate on 150mm LB-2% agar plates supplemented with carbenicillin. After incubation for 14 h at 37°C, colonies (∼900,000 total) were scraped from the agar plates into LB, pooled, and the plasmid DNA was extracted from ∼1×10^11^ cells using a midiprep kit. This pooled Mobile-CRISPRi library was transformed by electroporation into *E. coli* mating strain sJMP3049, plated at a density of ∼50,000 cells/plate on 150mm LB-2% agar plates supplemented with carbenicillin and DAP. After incubation for 14 h at 37°C, colonies (∼1,000,000 total) were scraped from the agar plates and pooled, the OD_600_ was normalized to 27 in LB with DAP and 15% glycerol and aliquots of the pooled CRISPRi library were stored as strain sJMP2942 at −80°C.

### Transfer of the Mobile-CRISPRi system to the *A. baumannii* chromosome

The MCi system was transferred to the *att*_Tn*7*_ site on the chromosome of *A. baumannii* by quad-parental conjugation of three donor strains—one with a mobilizable plasmid (pTn7C1) encoding Tn7 transposase, another with a conjugal helper plasmid (pEVS74), and a third with a mobilizable plasmid containing a Tn7 transposon encoding the CRISPRi system—and the recipient strain *A. baumannii* 19606. A detailed mating protocol for strains with individual sgRNAs was described previously (60). Briefly, 100 µl of culture of donor and recipient strains were added to 600 µl LB, pelleted at ∼8000 x g, washed twice with LB prior to depositing cells on a nitrocellulose filter (Millipore HAWP02500) on an LB plate, and incubated at 37°C, ∼5 hr. Cells were removed from the filter by vortexing in 200 µl LB, serially diluted, and grown with selection on LB-gent plates at 37°C.

For pooled library construction, Tn*7* transposase donor (sJMP2644), conjugation helper strain (sJMP2935), and recipient strain (sJMP490) were scraped from LB plates with appropriate selective additives into LB and the OD_600_ was normalized to ∼9. An aliquot of sJMP2942 pooled library strain was thawed and diluted to OD_600_ of ∼9. Eight ml of each strain was mixed and centrifuged at 8000xg, 10 min. Pelleted cells were resuspended in 4 ml LB, spread on two LB agar plates, and incubated for 5hr at 37°C prior to resuspension in LB + 15% glycerol and storage at −80°C. Aliquots were thawed and serial dilutions were plated on LB supplemented with gent (150) and LB. Efficiency of trans-conjugation (colony forming units on LB-gent vs. LB) was ∼1 in 10^7^. The remaining frozen stocks were plated on 150 mm LB plates solidified with 2% agar and supplemented with gent (150) and incubated for 16 h at 37°C. Cells were scraped from plates and resuspended in EZRDM (Teknova) + 25mM succinate + 15% glycerol at OD_600_ = 15 and aliquots were stored at −80°C as strain sJMP2949.

### Library growth experiment

The *A. baumannii* essential gene CRISPRi library (sJMP2949) was revived by dilution of 50 µl frozen stock (OD_600_ = 15) in 50 ml LB (starting OD_600_ = 0.015) and incubation in 250 ml flasks shaking at 37°C until OD_600_ = 0.2 (∼2.5 h) (timepoint = T0). This culture was diluted to OD_600_ = 0.02 in 4 ml LB with 1mM IPTG and antibiotics (colistin, imipenem, meropenem, rifampicin, and no antibiotic control) in 14 ml snap cap culture tubes (Corning 352059) in duplicate and incubated with shaking for 18 h at 37°C (T1). These cultures were serially diluted back to OD_600_ = 0.01 into fresh tubes containing the same media and incubated with shaking for 18 h at 37°C again (T2) for a total of ∼10-15 doublings. Cells were pelleted from 1 ml of culture in duplicate at each time point (T0, T1, T2) and stored at −20°C. Final antibiotic concentrations were (in µg/ml): colistin (Sigma C4461): 0.44 and 0.67, imipenem (Sigma I0160): 0.06 and 0.09, meropenem (Sigma 1392454): 0.11 and 0.17, and rifampicin (Sigma R3501): 0.34.

### Sequencing library samples

DNA was extracted from cell pellets with the DNeasy gDNA extraction kit (Qiagen) according to the manufacturer’s protocol, resuspending in a final volume of 100 µl with an average yield of ∼50 ng/µl. The sgRNA-encoding region was amplified using Q5 DNA polymerase (NEB) in a 100 µl reaction with 2 µl gDNA (∼100 ng) and primers oJMP697 and oJMP698 (nested primers with adapters for index PCR with Illumina TruSeq adapter) according to the manufacturer’s protocol using a BioRad C1000 thermocycler with the following program: 98°C, 30s then 16 cycles of: 98°C, 15s; 65°C, 15s; 72°C, 15s. PCR products were purified using the Monarch PCR and DNA Cleanup and eluted in a final volume of 20 µl for a final concentration of ∼20 ng/µl).

Samples were sequenced by the UW-Madison Biotech Center Next Generation Sequencing Core facility. Briefly, PCR products were amplified with nested primers containing i5 and i7 indexes and Illumina TruSeq adapters followed by bead cleanup, quantification, pooling and running on a Novaseq 6000 (150bp paired end reads).

### Library data analysis

For more information on digital resources and links to custom scripts, see Table S4.

### Counting sgRNA Sequences

Guides were counted using *seal.sh* script from the *bbtools* package (Release: March 28, 2018). Briefly, paired FASTQ files from amplicon sequencing were aligned in parallel to a reference file corresponding to the guides cloned into the library. Alignment was performed using *k*-mers of 20 nucleotide length—equal to the length of the guide sequence.

### Condition Comparisons – Quantification and Confidence

Log_2_-fold change and confidence intervals were computed using *edgeR*. Briefly, trended dispersion of guides was estimated and imputed into a quasi-likelihood negative binomial log-linear model. Changes in abundance and the corresponding false discovery rates were identified for each guide in each condition individually. Finally, log2-fold abundance changes were calculated by taking the median guide-level log2-fold change; confidence was calculated by computing the Stouffer’s *p*-value (*poolr R* package) using FDR for individual guides across genes.

### Knockdown-Response Curves

Code was adapted from the *drc* (*DoseResponse*) *R* package to generate 4-parameter logistic curves describing the relationship between predicted knockdown (independent) and the log_2_-fold change in strain representation (dependent) for all (∼10) mismatch guides per gene.

### Data Sharing Plan

Raw data will be deposited in the Sequence Read Archive (SRA), code used to analyze the data will be available on GitHub, and plasmids will be available from Addgene. Other reagents and protocols are available upon request.

## Acknowledgements

This work was supported by a Career Transition Award from the NIH National Institute of Allergy and Infectious Diseases (K22AI137122). R.D.W. was supported by the Predoctoral Training Program in Genetics (NIH 5T32GM007133). J.S.T. was supported by the Biotechnology Training Program (NIH 5T32GM135066) and a GRFP from the NSF. We thank ChatGPT for assistance in developing the CoMBaT-seq acronym. We thank Agilent Technologies for providing SurePrint Oligonucleotide libraries and Laura Whitman for oligo synthesis support and the University of Wisconsin Biotechnology Center for technical support with Illumina sequencing.

